# Aquaculture potential of Bohai Red and its hybrid with *Agropecten irradians concentricus* in southern China

**DOI:** 10.1101/2019.12.29.890160

**Authors:** Gaoyou Yao, Jianqiang Li, Yuyuan Wu, Xiaoying Su, Jie Tan, Zhigang Liu

**Affiliations:** Engineering Research Center of Healthy Breeding for Economic Invertebrates in the South China Sea,Zhanjiang, Guangdong,524088,China; College of Fisheries, Guangdong Ocean University, Zhanjiang 524025, China

**Author notes:** Corresponding author: College of Fisheries, Guangdong Ocean University, Zhanjiang 524088, PR China. *E-mail address.

**Keywords:** *Argopecten irradians concentricus*, Bohai Red, Hybridization, Temperature tolerance, Heterosis

## Abstract

*Argopecten irradians concentricus* (Say), one of four geographic subspecies of the bay scallop, has become the major cultured species in southern China since its introduction in 1995. However, its population has been significantly reduced due to high mortality.Also, poor growth rate has been observed following decades of culture.Therefore, the introduction or creation of new varieties is an urgent need. This study describes the first introduction of the new strain, Bohai Red, from the north to southern China. Hybridization trials were conducted between the recently introduced new strain and the local specie, *A. irradians concentricus* (Say). The success of hybridization was confirmed by SSR maker.The adult and juvenile Bohai Red cannot tolerate high temperatures compared to *A. irradians concentricus* (Say), whether in natural waters or under laboratory conditions. Fertilization rate for *A. irradians concentricus* (Say) × Bohai Red exceeded 80%, and hatching rate was 70%. The hybrids exhibit heterosis in survival rate, growth rate, and high-temperature tolerance, demonstrating substantial potential to replace *A. irradians concentricus* (Say) in southern China.

## 1. Introduction

As an economically important marine bivalve, the bay scallop *Argopecten irradians* (*A.i.*) has four geographic subspecies: *A.i.irradians* (Lamarck),*A.i.amplicostatus* (Dall), *A.i. concentricus* (Say), and *A.i.taylorae* (Petuch), which are naturally distributed along the coast of the Atlantic Ocean and the Gulf of Mexico (Wilbur, Gaffney, 1997). The original stocks of *A. irradians concentricus* (Say) were first introduced to China from North Carolina in 1995 (Zhang et al, 2000). Due to the fast growth rate, and large adductor muscle, the production of the bay scallop reached about 600,000 tons in 2007 (Guo, 2009; Wang et al, 2014). However, since the Say scallop is a hermaphroditic animal, significant decreases in population due to high mortality, and also poor growth rate had been noticed following decades of culture. (Wang et al, 2014; Wang et al, 2007; Zheng et al, 2008). Efforts such as intra-specific and inter-specific hybridization did result in improved growth rate, but its tolerance to environmental conditions such as *Argopecten purpuratus×A. irradians concentricus, Argopecten irradians×A. irradians concentricus and Mimachlamys nobilis×A. irradians concentricus* were minimal.New varieties to replace bay scallops in southern China have not been identified and the problem of germplasm degradation in the southern bay scallop has not been effectively solved.

Bohai Red is a new scallop strain formed from the hybridization of the *Argopecten purpuratus* and bay scallop *Argopecten irradians* (Lamarck) through six generations of selection which exhibits excellent heterosis in growth. Compared to bay scallops, this strain is 125.9–138.9% larger by whole body weight and has a 145.4–156.2% larger adductor muscle. So the *A. i. irradians* (Lamarck) was quickly being replaced by the Bohai Red after the new strain passed the certification of Fisheries Law of People’s Republic of China. By 2016, in the north of China, Bohai Red production reached 556470 tons, replacing more than 50% of the traditional culture areas of the *A. i. irradians* (Lamarck) (Wang et al, 2017; Wang et al, 2011). But as a northern cultured scallop variety, it is only been cultured in Bohai Bay of north china but not in the south where the culture area is 2-3 times higher than that of the northern culture area. Considering the degradation of *Argopecten irradians concentricus* in south China, Bohai Red may provide an opportunity for genetic improvement of *A.irradians concentricus.* However, it is unknown if Bohai Red can be successfully introduced in the south due to the significant temperature differences between the north and south.

Although the mechanism of heterosis still remains enigmatic (Wang et al, 2017; Wang et al, 2011), hybridization is considered as one of the most effective methods for genetic improvement in livestock and plant production. In aquaculture, hybridization has been extensively applied to enhance growth rate, survival rate, and environmental tolerance in the oyster (Hedgecock et al, 1995; Kong et al, 2017), abalone (You et al, 2009; You et al, 2015) and scallop (Cruz, Ibarra, 1997; Cruz et al, 1998; Wang et al, 2011). For example, due to high-temperature tolerance and accelerated growth rate, the Penglai red hybrid between *Chlamys farreri* and *Patinopecten yessoensis* was commercially introduced in China in 2003 (Sun et al, 2004; Yang et al, 2004). Since Bohai Red is the hybrid between the *Argopecten purpuratus* and *Argopecten irradians* (Lamarck), it is likely that Bohai Red and *A. irradians concentricus* (Say) can be successfully crossed to produce offspring with heterosis, furthermore, we are interested in the ability of hybrids to tolerate the high temperatures as this is an obligatory adaptation for southern China.

In order to improve the sustainability of bay scallops in China, the Bohai Red, which was first introduced from the north of China, was evaluated for growth and survival rate in southern China to determine if it will be able to adapt to the environment of southern China. In addition, crossbreeding between the Bohai Red and *Argopecten irradians concentricus* was conducted and heteroses in growth and survival traits were assessed by comparing the hybrid crosses with purebred offspring.

## 2. Material and methods

### 2.1. Comparisons of survival in two broodstocks

One thousand and one hundred adult Bohai Red scallops, which were 40 mm–50 mm in shell length, were collected in Qindao, Shangdong, China (35.7°N,120.0°E) by professor Chunde Wang, who created the new strain. Five hundred adult *Argopecten irradians concentricus* were collected from culture stocks in Beibu bay (20.4°N, 109.9°E) with shell length ranging from 40 to 50 mm. The two stocks were cultured in Beibu bay, Guangdong, China (20.8°N, 109.7°E) from December to April. The water temperature was measured at 16:00 each day to calculate the monthly average water temperature. Dead scallops were collected to calculate the survival rate.

### 2.2. Gamete collection and hybridization

When the broodstocks of Bohai Red (designated as Z) and *Argopecten irradians concentricus* (designated as M) were sexually mature, the two broodstocks were hybridized. Ten sexually mature individuals were chosen from each stock to spawn by flowing water stimulation for two hours followed by injection of 0.1 mL of 0.05 mmol/L serotonin (5-hydroxytryptamine, Sigma, USA) into the adductor muscle (Cruz, Ibarra, 1997). All scallops begin to release eggs after 30 minutes of releasing of the sperm. At this time, all scallops were separately placed in a 2-L box filled with seawater of the same temperature to collect eggs or sperm. The eggs were discarded once the first polar bodies were identified under the microscope following 15 minutes of egg collection. Eggs and sperm of the two stocks were collected separately. The gametes from different broodstock were mixed with eggs from another stock to create hybrids groups (M♀×Z♂, M♂×Z♀). Those from the same stock were used to create parental groups (M♀×M♂, Z♀×Z♂). Three parallel parental and hybrid groups were created, and a limited number of fertilized eggs were collected to evaluate fertilization success and survival to D-stage larvae.

### 2.3 Larvae and juvenile rearing

The larvae were kept in 500-L aerated plastic tanks at 22–25°C. The *Isochrysis zhanjiangensis* and *Chlorella vulgaris* were mixed in equal amounts and fed. The density of larvae was maintained at 20 larvae/ml. The collectors were placed in the nursery tanks when 30% of larvae had eyespots. Three 2-ml samples were taken from each group and the numbers of larvae were used to calculate the density. The survival rate was recorded based on the live larvae on different days. The larvae were removed from the collector when the shell length of larvae reach 450 um and was placed in a 0.15mm mesh screen to transfer them to the Beibu Bay, China,(20.4°N,109.9°E) for the nursery stage. The spats were sorted at a day intervals, first into larger mesh bags and finally into adult cages. Forty progenies per replicate were randomly sampled and their growth index (shell length/mm, shell height/mm and weight/g) was measured with vernier calipers (0.02 mm accuracy) and an electronic balance (0.01 g accuracy) every month(from seeding to harvesting for about 200d).

### 2.4 Genetic confirmation

DNA was extracted from samples of the hybrids and parents progeny using the marine animal DNA extraction kit (transgene, China). The SSR primer information reference literature (Jie et al, 2018).The procedure was as follows: (1) Primer M13 (containing HEX HEX or 6-FAMFAM fluorescent material, sequence 5’ -CACGACGTTGTAAAACGACCACGACGTTGTAAAACGAC) was added to the 5’ end of the forward primer (completed by Sangon Biotech, China), and the reverse primer was unchanged. (2) PCR reactions involving genomic DNA templates from the parents and hybrids. The PCR was conducted as follow: 3 min at 95°C (pre-denaturation); 10 cycles of denaturing for 30 s at 95°C, annealing for 30 s at 60°C and extending for 30 s at 72°C;20 cycles of denaturing for 30 s at 95°C, annealing for 30 s at 55°C and extending for 30 s at 72°C;followed by a final extension of 6 min at 72°C(3) 50 pg of PCR product and with 10 μl of Buffer (HiDi formamide and ROX500 in a ratio of 99:1) were mixed, then rapidly cool on ice-water mixture immediately after 5 min at 98°C. The capillary electrophoresis was performed on the 3730XL (Applied Biosystems) and the results were analyzed using Genemapper software.

### 2.5 Temperature tolerance test

To determine the temperature tolerance, hybrid cohorts and inbred cohorts weighing 0.22–0.24 g, with shell lengths of 10.8–10.9 mm, were first acclimated to the laboratory conditions for 24 h prior to the test. The temperature groups were set at 18°C and 30°C in 10-L plastic containers equipped with temperature control units. Sixty individuals were randomly selected from each cohort, and three replicates were assigned. The scallops were fed with *Chlorella vulgaris* twice a day. The seawater was changed and the dead scallops were picked out to calculate the survival rate every 24 h. The final shell length was measured to calculate the growth rate after 15 d.

### 2.6. Heterosis estimate and data analysis

The data are expressed as mean±SD. Statistical analyses were performed using IBM SPSS, Version 19 (IBM SPSS Inc, Chicago, IL, USA). The differences in survival and growth among the experimental lines and heterosis were analyzed using one-way ANOVA, and significance for all analyses was set at *P* < 0.05. Mid-parent heterosis was calculated using the formula (Falconer, 1996).

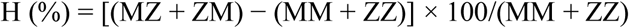

where MZ and ZM are the mean survival rate (or weight, size) of the hybrid F1 cohort, and MM, ZZ is the mean survival rate (or weight, size) of the *A. irradians concentricus* Say and Bohai Red, respectively.

Single-parent heterosis was calculated using the formula (Cruz, Ibarra, 1997):

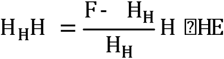

where H_X_ is the single-parent heterosis of stock X, P_X_ is the mean size (or weight, survival rate) of the cohort parental stock X (ZZ, MM), and F is the mean size (or weigh survival rate) of the hybrid cohort (MZ, ZM).

## 3. Result

### 3.1 Comparisons the survival of two broodstocks in the southern waters

The water temperature in the culturing area was measured continuously over five months. The lowest average temperature, 18°C, was observed in February, while the highest average temperature was observed in April at 26.8°C. The two broodstocks exhibited different survival rates at the end of the 5 months of growth and the survival rate for Bohai Red was closely related to temperature (Fig. 1). Survival rate exceeded 90% for two broodstocks from December to March, but the survival rate dropped to 70.2% for Bohai Red when the average water temperature reached 26.8°C in April while the survival rate of *Argopecten irradians concentricu* remained high survival rate(92%) during this period.

**Fig. 1.**
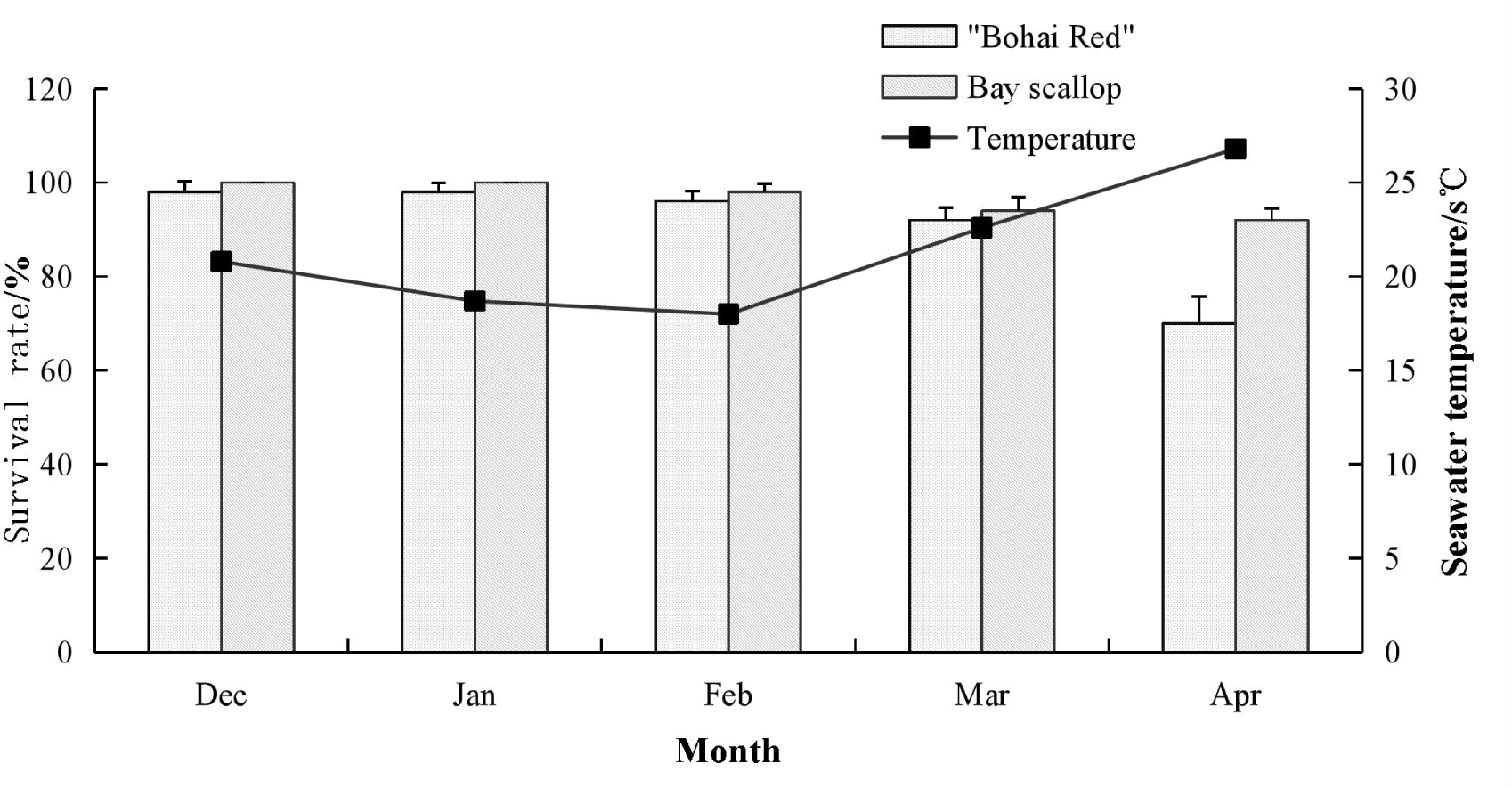
Survival rate of Bohai Red and Bay Scallop from December to April

### 3.2 Fertilization and hatching rates

The hatching and fertilization rates are displayed in Table 2. The highest hatching rate was recorded in MM (90.63%) with the highest fertilization in MZ (78.80%), While ZM had the lowest hatching (88.31%) and fertilization rate (74.6%).There was no significant difference between all groups (*P >* 0.05).

**Table 1.**
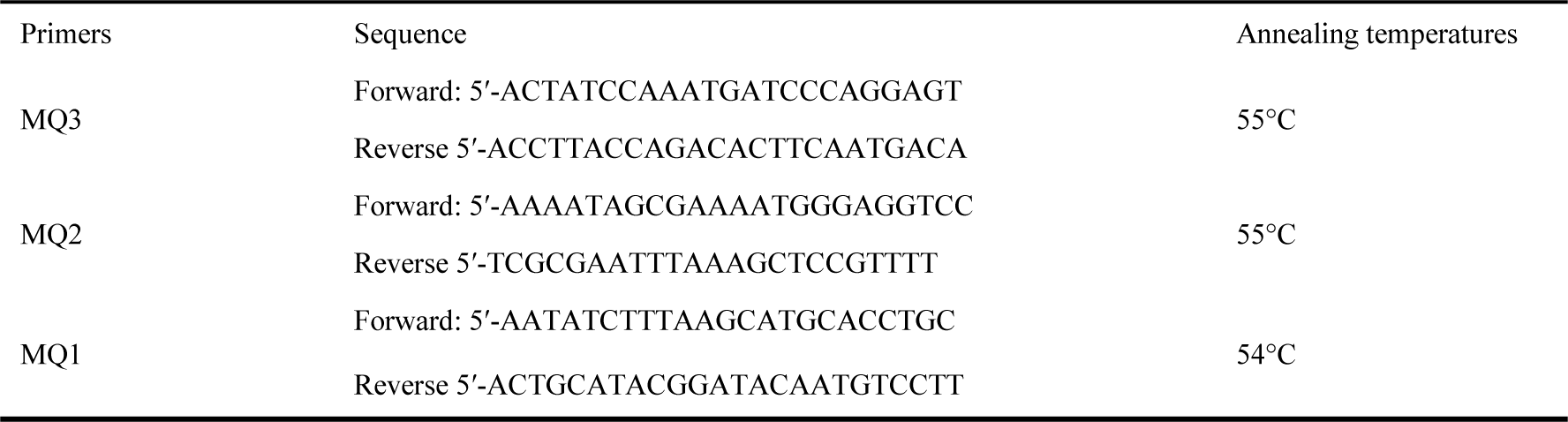
Primers for parentage analyses

**Table 2.** Fertilization and hatching rates in different groups

|  | MM | MZ | ZZ | ZM |
| --- | --- | --- | --- | --- |
| Fertilization rate (%) | 90.63±0.69<sup>a</sup> | 89.41±1.20<sup>a</sup> | 88.73±0.87<sup>a</sup> | 88.31±1.77<sup>a</sup> |
| Hatching rate (%) | 77.40±5.77<sup>a</sup> | 78.80±3.90<sup>a</sup> | 76.40±5.94<sup>a</sup> | 74.60±4.04<sup>a</sup> |

Notes: The same letters indicate no significant difference ( $P > 0.05$ )

### 3.3 Growth and survival during the larval stage

During the larval stage, there were significant differences between the four groups in shell length (Table 3). The hybrid group (MZ, ZM) exhibited substantial heterosis (4.50%–18.57%) during the larval stage which gradually increased over time. The single parent heterosis of MZ was significantly higher than that of the ZM group at 5, 10, 20 and 30 d. The growth rates of the two-hybrid groups were greater than the parent groups (MM ZZ), with middle-parental heterosis of 21.59%. Shell length of MZ was significantly greater than all other groups, while no significant difference was found between the pure groups (*P* > 0.05). In term of survival rate, it ranged from 41.83–49.09% with no significant difference between all groups at the larval stage.

**Table 3.** Growth and survival of reciprocal and parental crosses at the larval stage

| Combination | Shell length (um) |  |  |  | Growth rate<br>(um/d) | Survival rate (%) |
| --- | --- | --- | --- | --- | --- | --- |
|  | 5 d | 10 d | 20 d | 30 d |  |  |
| MM | 112.63±8.93<sup>b</sup> | 148.30±10.50<sup>b</sup> | 300.57±12.49<sup>b</sup> | 625.13±35.28<sup>c</sup> | 20.50±1.16<sup>c</sup> | 41.83±3.41<sup>a</sup> |
| MZ | 116.33±10.14<sup>a</sup> | 153.13±9.66<sup>a</sup> | 353.30±21.16<sup>a</sup> | 754.60±83.23<sup>a</sup> | 25.53±3.06<sup>a</sup> | 49.09±5.73<sup>a</sup> |
| ZZ | 103.47±8.11<sup>d</sup> | 140.23±7.38<sup>c</sup> | 290.80±10.46<sup>c</sup> | 603.13±45.34<sup>c</sup> | 19.98±1.61<sup>c</sup> | 47.87±4.35<sup>a</sup> |
| ZM | 109.50±8.13<sup>c</sup> | 146.73±7.04<sup>b</sup> | 313.63±24.25<sup>b</sup> | 701.73±64.18<sup>b</sup> | 23.69±2.39<sup>b</sup> | 47.41±4.99<sup>a</sup> |
| H<sub>M</sub>(%) | 3.28 | 3.26 | 17.54 | 20.70 | 24.54 | 17.36 |
| H<sub>Z</sub>(%) | 5.83 | 4.64 | 7.80 | 16.35 | 18.57 | -0.96 |
| H (%) | 4.50 | 3.93 | 12.78 | 18.57 | 21.59 | 7.58 |

Notes: The same letters indicate no significant difference ( $P > 0.05$ ).

**Table 4.** Comparisons of growth and survival of the hybrid and inbred cohorts at harvest

| Combination | Shell length (mm) | Growth rate (mm/d) | Weight (g) | Growth rate (g/d) | Survival rate (%) |
| --- | --- | --- | --- | --- | --- |
| MM | 47.76±4.10<sup>c</sup> | 0.40±0.034<sup>c</sup> | 21.87±4.17<sup>c</sup> | 0.18±0.035<sup>c</sup> | 81.60±3.04<sup>b</sup> |
| MZ | 53.64±5.03<sup>a</sup> | 0.45±0.042<sup>a</sup> | 28.39±6.62<sup>a</sup> | 0.24±0.055<sup>a</sup> | 87.52±3.00<sup>a</sup> |
| ZZ | 50.93±3.23<sup>b</sup> | 0.42±0.027<sup>b</sup> | 24.52±6.87<sup>b</sup> | 0.20±0.057<sup>b</sup> | 44.46±3.33<sup>c</sup> |
| ZM | 51.47±4.95<sup>b</sup> | 0.43±0.041<sup>b</sup> | 24.70±4.90<sup>b</sup> | 0.21±0.041<sup>b</sup> | 85.62±2.37<sup>ab</sup> |
| H<sub>M</sub>(%) | 12.31 | - | 29.81 | - | 7.25 |
| H<sub>Z</sub>(%) | 1.05 | - | 0.73 | - | 92.58 |
| H | 6.51 | - | 14.44 | - | 37.35 |

### 3.3 Growth and survival during growth-out stage

The survival and growth rates of the inbred and hybrid cohorts are shown in Fig. 2 and Fig. 3. The growth rates for shell length of the hybrid groups (MZ, ZM) were higher than those of the parental crosses (MM, ZZ), and the shell length for MZ reached 53.64 mm with a growth rate 0.45 mm/d, the greatest rate among the hybrid groups (H_M_ = 37.20%, H_Z_ = 27.14% and H = 32.23%). The shell length for the MM group was significantly lower than the other groups at 47.76 mm (*P* > 0.05). The middle-parental heterosis of the whole body weight was greater than shell length heterosis, with an average value of 14.41%. While the whole body weight growth rates for the hybrid groups (MZ, ZM) were greater than the parental crosses (MM, ZZ), there was no significant difference between ZM (24.70 g) and ZZ (24.52 g). The average values were 0.73% for H_Z_ and 29.81% for H_M,_and 14.44 %for H.

**Fig. 2.**
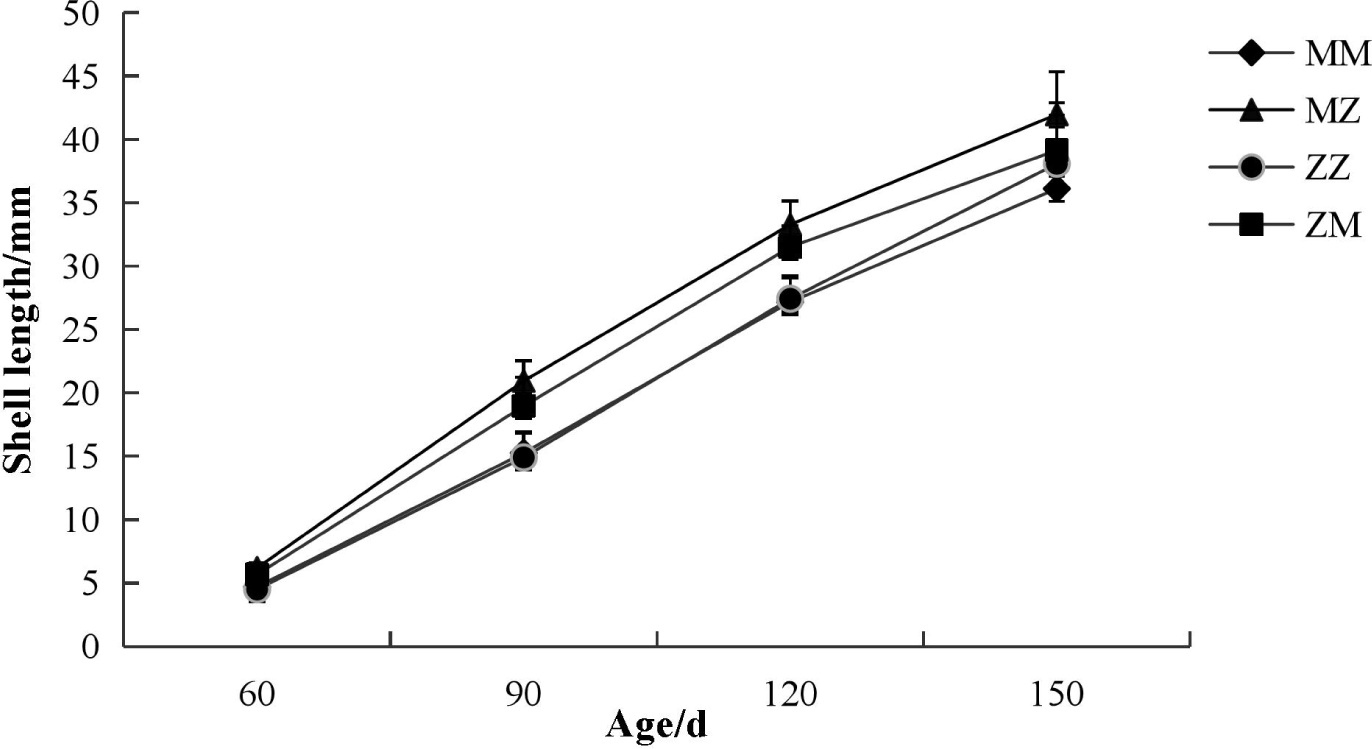
Shell length of inbred and hybrid scallops from 60 to 150 day age

**Fig. 3.**
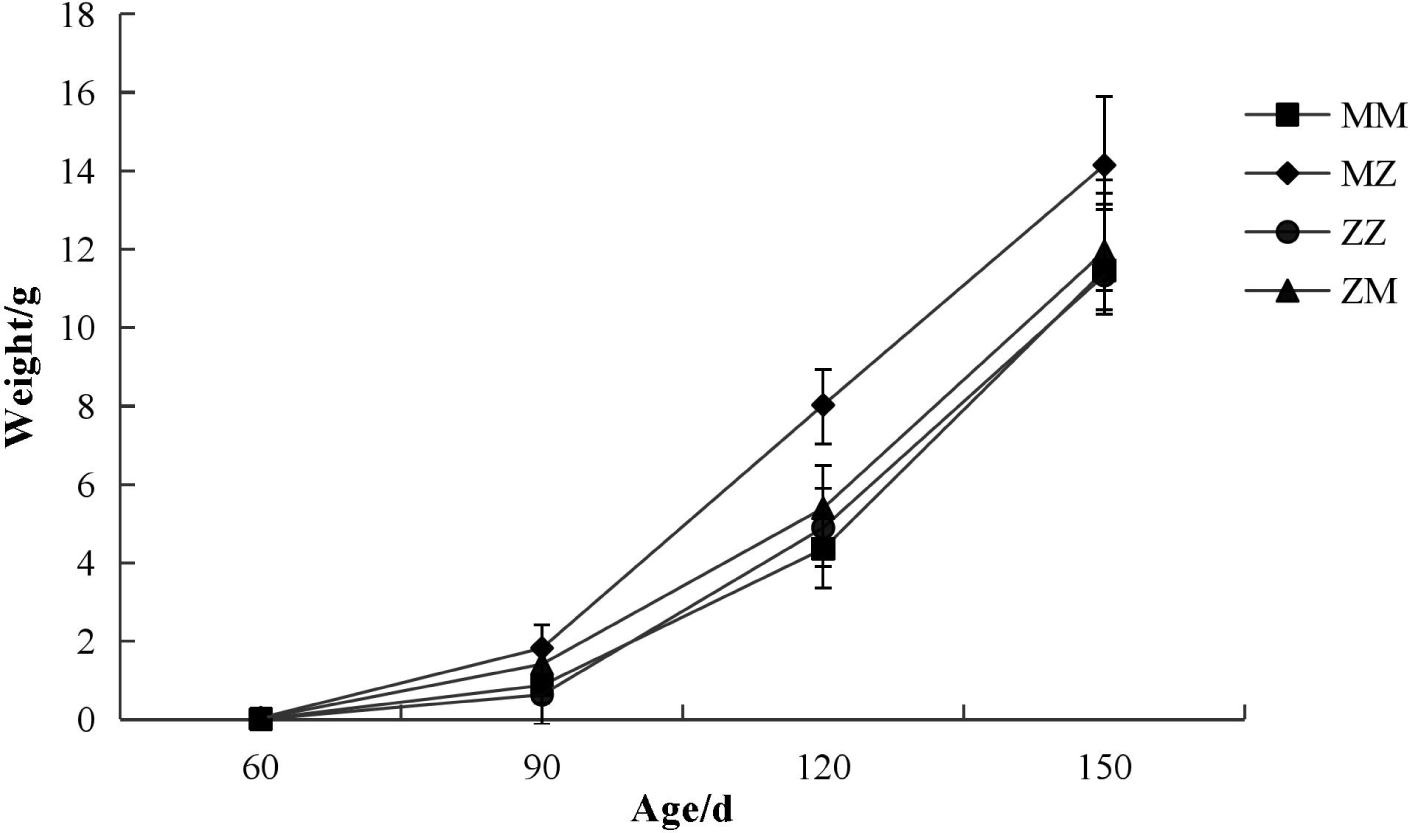
Body weight of inbred and hybrid scallops from 60 to 150 day age.

In terms of survival rate, mortality rose rapidly for the ZZ group at high water temperatures. The survival rate for the ZZ group was only 44.46% at harvest, while the other groups were all higher than 80%. No significant differences in survival rates were observed between hybrid cohorts (*P* > 0.05), with average values of 7.25% for H_M_, 92.58% for H_Z_ and 27.35% for H.

### 3.4 Parentage analyses

Parentage analyses with the three SSR primers (MQ1, MQ2, MQ3) are shown in Fig.4 For the *Argopecten irradians concentricus* Say, only one band ~146 bp was amplified by MQ1, one band ~132 bp was amplified by MQ2 and two bands, ~116 bp, and ~124 bp, by MQ3. For the Bohai Red only one band was amplified by MQ1(~148 bp),MQ2(~134bp) and MQ3(~134bp)In all the hybrids, only two bands were amplified, one was only appeared in the *Argopecten irradians concentricus* (Say) and another only appeared in Bohai Red, indicating that the success of hybridization can be confirmed.

**Fig. 4.**
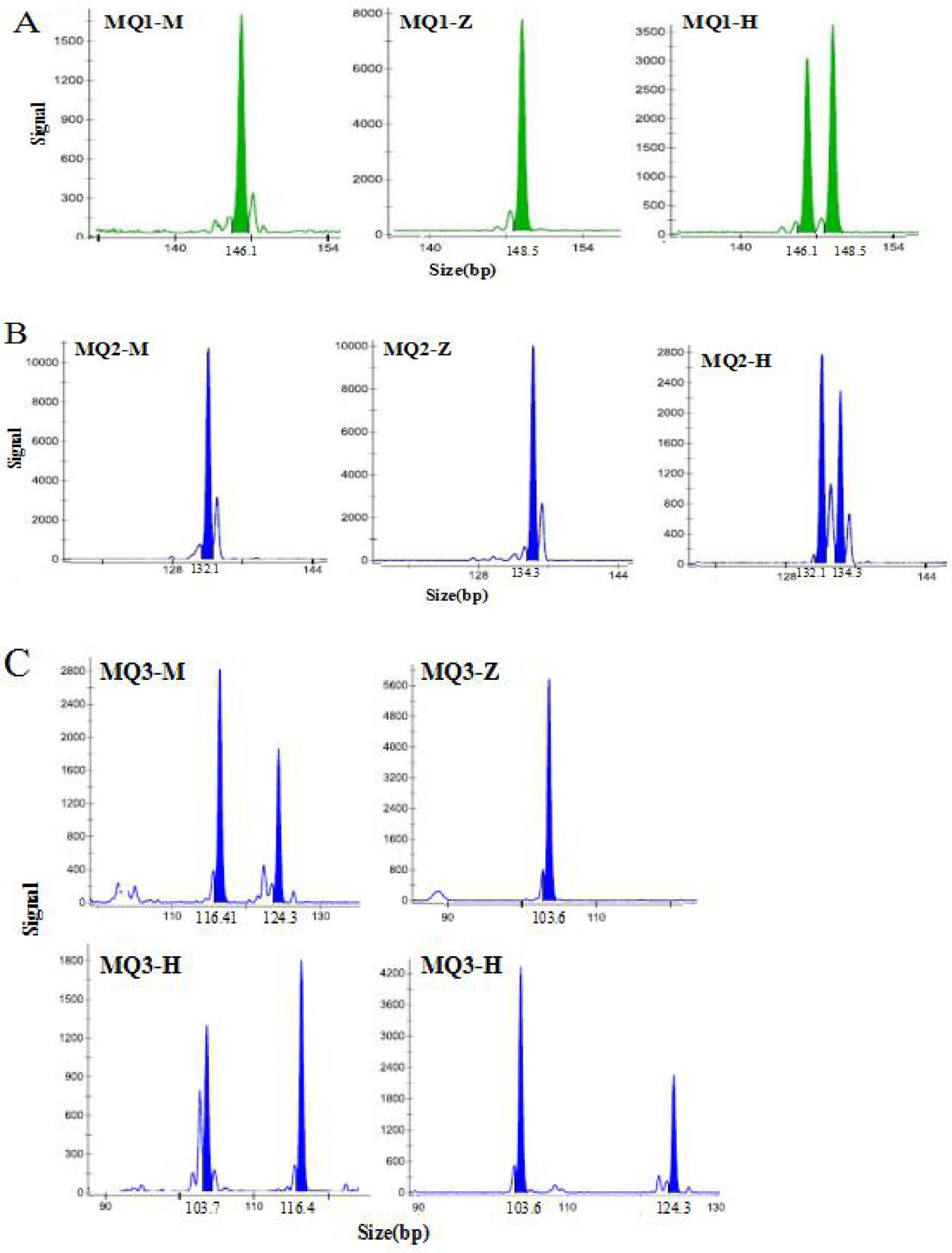
Genotyping results of three SSR loci in “Bohai Red”, *A.i.concentricus* and their hybrid. Note: MQ-M: Genotyping results of *A.i.concentricus* MQ-Z: Genotyping results of “Bohai Red” MQ-H: Genotyping results of hybrid

### 3.5 Temperature tolerance test

Average summer temperature (30°C) and average winter temperature (18°C) in southern China were used for temperature tolerance test.The growth and survival rates of the hybrids (MZ, ZM) and inbreds (MM, ZZ) under laboratory conditions are presented in Figs.5 and Figs.6. The results indicate that the high-temperature tolerance ability of MM was greatest with growth rate (0.3084 mm/d) and survival rate(78.34%),followed by ZM and MZ.The ZZ was weakest with growth rate (0.1015mm/d) and survival rate(25%).There was no significant difference recorded among the groups in the low-temperature(18°C). The inbred ZZ exhibited the highest growth rate 0.30 mm/d (P>0.05) in the low-temperature environment while the worse growth rate was recorded under high temperature (30°C) tolerance with survival rate 25% and growth rate 0.1015mm/d (*P*<0.05).Our result demonstrate that the ability to tolerate the high temperature of hybrids was improved compared to Bohai Red.

**Fig. 5.**
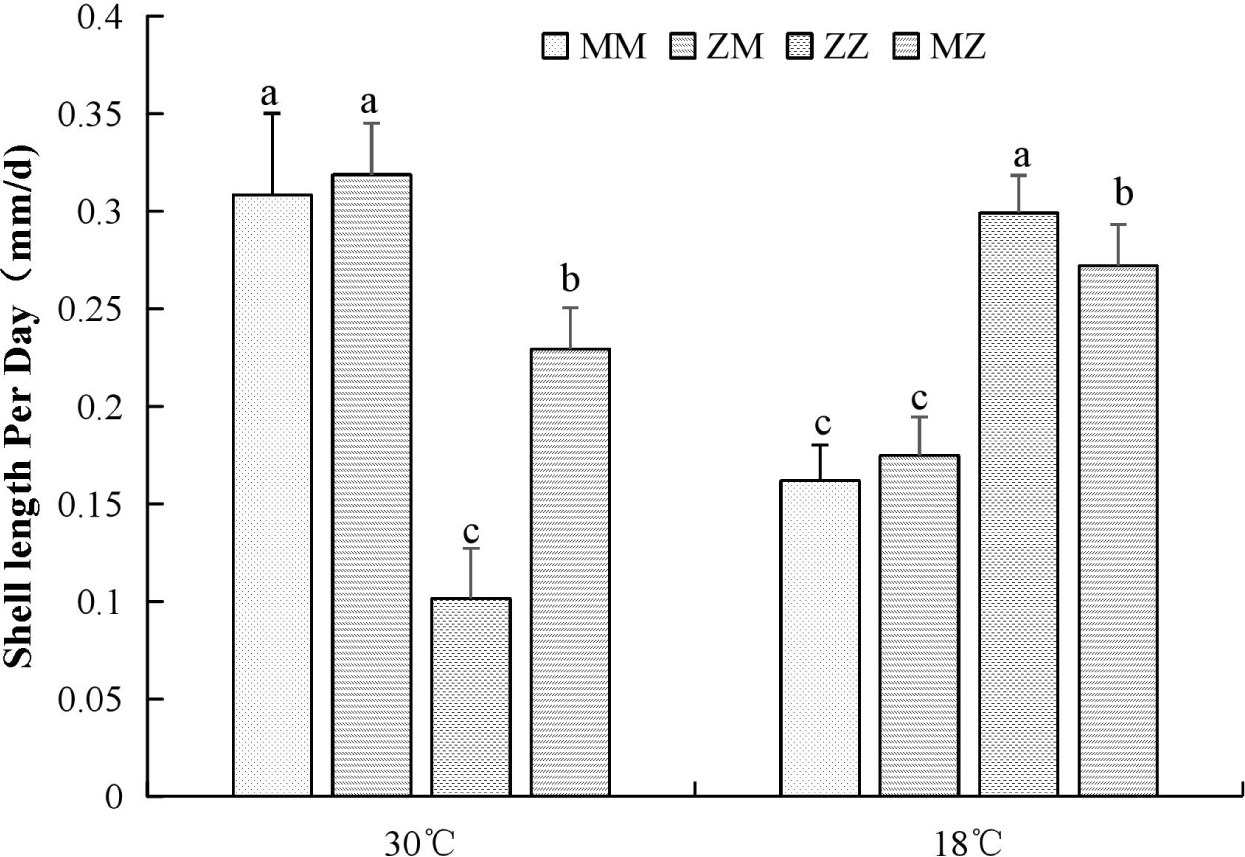
The growth of hybrid and inbred cohorts under different water temperatures in the laboratory.

**Fig. 6.**
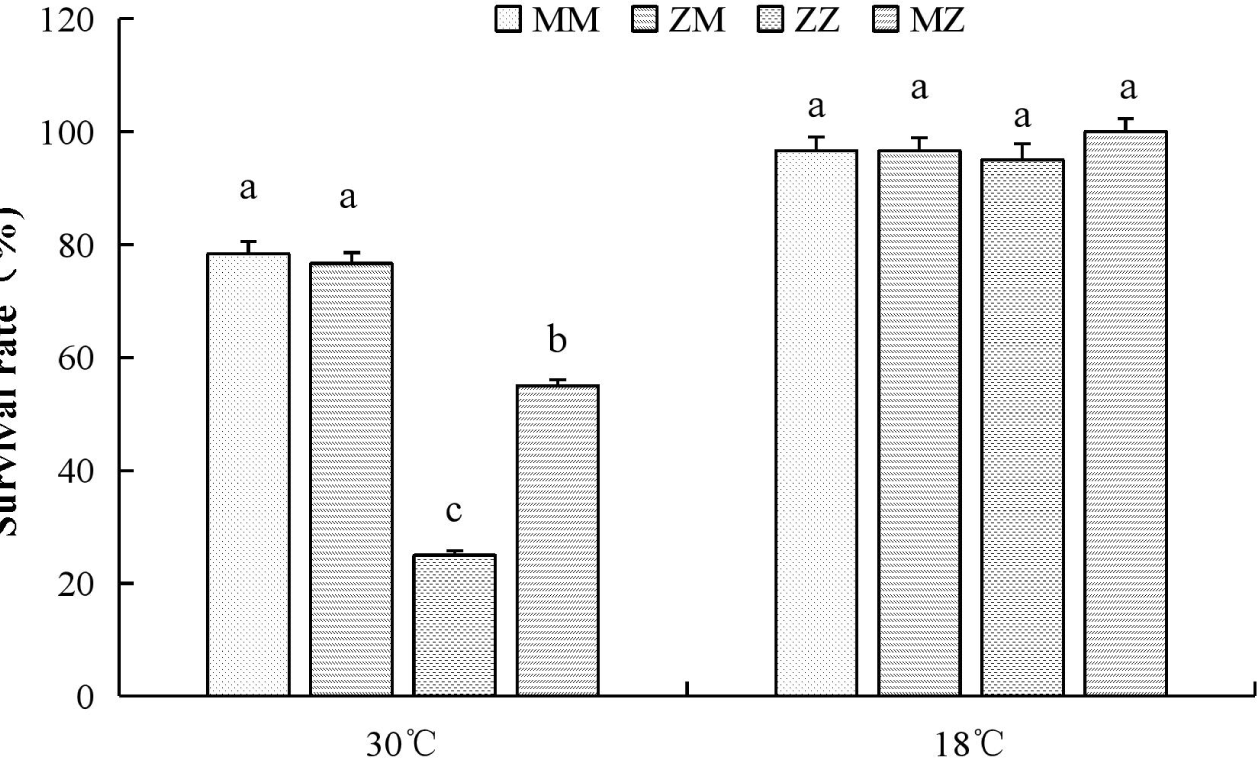
Survival rate of hybrid and inbred cohorts under different water temperatures in the laboratory.

## 4 Discussion

In this study, Bohai Red was introduced into the south of China for the first time.We found that the adult and juvenile Bohai Red cannot tolerate high temperatures compared to *A. irradians concentricus*, whether in natural waters or under laboratory conditions.The results matched the reports about different ability of temperature tolerance among the two species.Preliminary results showed that the critical thermal maxima were 32°Cfor *Argopecten irradians concentricus*(Liu, 2007), compared to 29 °C for Bohai Red(Wang et al, 2017).High mortality was observed in Bohai Red with the increasing temperature demonstrating that thermal stress may also be the primary challenge for the introduction of Bohai Red, and it may not be suitable for direct introduction to southern China where the maximum water temperature reaches 32°C in summer. However, the hybrids between *A. irradians concentricus and* Bohai Red are adaptable to south water which suggests that ability of hybrids to tolerate the high temperatures was increased through hybridization.Same results had appeared in *Haliotis discus hannai* ×*H. fulgens*(You et al, 2015), red abalone♀ × Pacific abalone♂(Cruz, Gallardo-Escárate, 2011)and *Argopecten purpuratus*×*Argopecten irradians* (Cruz, Gallardo-Escárate, 2011). The hybrids could change physiological status to get more resistant to disease and thermal stresses than their parents(Liang et al, 2014; Nan et al, 2016).

Summer mortality is multifactorial process, including genetics, pathology, physiology, and immunology (Samain et al, 2008) which could be caused by an interaction between abiotic environmental factors and biotic at high water temperatures(Stone et al, 2014).The reasons for high mortality of Bohai Red in the high temperature were complex, while the temperature is only one of the factors related to mortality.our results demonstrate that hybridization could be an effective way to improve the ability of Bohai Red to tolerate high temperature conditions.

Bohai Red, non-native genetic material, was used for genetic improvement of native species (*A.irradians concentricus*) through interspecies hybridization.Compared to *A. irradians concentricus* (Say), shell lengtht and whole body weight of the hybrid increased by 12.31%, and 29.81%, respectively. Generally, hybrids could develop heterosis in disease resistance, growth, or other traits than parental species through hybridization that promotes genomic transfer between species and produces phenotypic and genotypic changes (Stone et al, 2014)(Stelkens et al, 2009; Zhang et al, 2014). Similarly, our results also have shown that the growth of the hybrids was likely to be greatly improved by the introgression gene flow between two parental species which it is likely an important plastic force in the morphological and genetic diversity of the hybrids(Bierne et al, 2006).In our study, the growth rate of MZ significantly exceeded that of ZM in hybrid groups.Differences between reciprocal crosses are also commonly observed in shellfish such as *Crassostrea hongkongensis*(Bierne et al, 2006)*, Argopecten irradians*(Zhang et al, 2014), it may be due to the cytoplasmic inheritance, sex-linked genes, or parental effects(Zhang et al, 2014).This study demonstrates that the cross between Bohai Red and *A. irradians concentricus* may be a good way in increasing growth of *A. irradians concentricus*.

Bay scallop is hermaphroditic animal that spawns sperm and eggs simultaneously, So it is hard to separate sperm and eggs at a large spawning scale, as performed in one-to-one crossbreeding (Zhang et al, 2007). Furthermore, there are phenomena of gynogenesis, natural polyploid, androgenesis in shellfish(Zhang et al, 2007), we can not confirm hybrids just basing on morphologies such as number of the ribs or shell colar (Hedgecock et al, 1995).Genetic identification of interspecific hybrids among shellfish has been reported using karyological studies, SSR(Hedgecock et al, 1995), CO I, 16S(Ahmed et al, 2008; An et al, 2005) and ITS(Xu et al, 2019).In our research, primes MQ1, MQ2 and MQ3 were developed as species-specific marker to distinguish *A. irradians concentricus* and “Bohai Red” as only one band ~146 bp was amplified by MQ1, only one band ~132 bp was amplified by MQ2 in *Argopecten irradians concentricu*s and only one band was amplified by MQ1(~148 bp),MQ2(~134bp) and MQ3(~134bp) in “Bohai Red”, but Both hybrids, two bands were amplified at the same time which can confirm the hybrid at the genetic level.

Three-way crosses were defined as: the hybridization of a binary hybrid of two varieties (lines) to another cultivar (line) with excellent traits(Yan et al, 2010).It can combine the ideal traits of three species to make the most of heterosis and had been widely used in the breeding of crops (Padhi et al, 2016) and livestock(Yamak et al, 2014). For shellfish in aquaculture, three-way crosses had only been reported in *Ruditapes philippinarum* (Yan et al, 2010).In this study, three-way crosses were conducted between *Argopecten irradians concentricus, Argopecten purpuratus, and Argopecten irradians irradians.* The results shown that compared to the heterosis which only in growth but not high temperature tolerance in hybridization of *Argopecten irradians concentricus*×*Argopecten irradians irradians*. *Argopecten irradians concentricus*×*Chlamys nobilis and Argopecten irradians concentricus*×*Argopecten purpuratus* (Nan et al, 2012),the ternary hybrid has shown heterosis in both growth and high-temperature tolerance which demonstrating that three-way crosses are also an effective method for genetic improvement in *A. irradians concentricus.* Furthermore, Shellfish usually spawn a lot(Wang, Li, 2010),for example, a bay scallop can produce between 500,000 and 600,000 gametes at a time (Lautert et al, 2012)and some of shellfish species are hermaphroditic animal which means that the inheritance of some ideal traits can be stabilized quickly, So we believe three-way crosses could suitable for shellfish breeding and have advantages overcrop and livestock in some aspects.

Although many experimental species of scallop hybridization have shown promising results and positive heterosis in aquaculture, only a few hybrids were commercially introduced until now.In the present study, we were excited to find that the hybrids of*A. irradians concentricus* and Bohai Red have improved in growth and tolerance to higher temperature when compared to parents. In order to produce the hybrid with stable inheritance and better growth performance in the future prospective research will focus on backcrossing with parental species and selective breeding to find strategies to maintain hybrid advantage.

## Acknowledgements

This work was sponsored by Key Research and Development Program of Guangdong (2019B020238002) and 2019 Innovation Team of Modern Agricultural in Guangdong(2019KJ146)]. The author would like to thank Chunde Wang for supporting the project. The authors are also grateful to Balu Liao for their assistance during the hatchery and adult grow-out stages.

## References

Ahmed, F., Koike, Y., Strüssmann, C.A., Yamasaki, I., Yokota, M., Watanabe, S., 2008. Genetic characterization and gonad development of artificially produced interspecific hybrids of the abalones, *Haliotis discus discus Reeve*, *Haliotis gigantea Gmelin* and *Haliotis madaka Habe*. Aquac Res 39, 532–541.

An, H.S., Jee, Y.J., Min, K.S., Kim, B.L., Han, S.J., 2005. Phylogenetic analysis of six species of pacific abalone (*Haliotidae*) based on DNA sequences of 16s rRNA and cytochrome c oxidase subunit I mitochondrial genes. Mar Biotechnol 7, 373–380.

Barton, N.H., 2001. The role of hybridization in evolution. Mol Ecol 10, 551–568.

Bierne, N., Bonhomme, F., Boudry, P., Szulkin, M., David, P., 2006. Fitness landscapes support the dominance theory of post-zygotic isolation in the mussels Mytilus edulis and *M. galloprovincialis*. Royal Society Proceedings B Biological Sciences 273, 1253–1260.

Cruz, F.L.D.L., Gallardo-Escárate, C., 2011. Intraspecies and interspecies hybrids in Haliotis: Natural and experimental evidence and its impact on abalone aquaculture. Rev Aquacult 3, 74–99.

Cruz, P., Ibarra, A.M., 1997. Larval growth and survival of two catarina scallop (*Argopecten circularis*, Sowerby, 1835) populations and their reciprocal crosses. J Exp Mar Biol Ecol 212, 95–110.

Cruz, P., Ramirez, J.L., Garcia, G.A., Ibarra, A.M., 1998. Genetic differences between two populations of catarina scallop (*Argopecten ventricosus*) for adaptations for growth and survival in a stressful environment. Aquaculture 166, 321–335.

Devanand, P.S., Rangaswamy, M., Ikehashi, H., 2000. Identification of hybrid sterility gene loci in two cytoplasmic male sterile lines in rice. Crop Sci 40, 640–646.

Falconer, D.S.M.T., 1996. Introduction to Quantitative Genetics. Longman.

Gaffney, P.M., Allen, S.K., 1993. Hybridization among Crassostrea species: a review. Aquaculture 116, 1–13.

Griffing, B., 1990. Use of a controlled-nutrient experiment to test heterosis hypotheses. Genetics 126, 753.

Guo, X., 2009. Use and exchange of genetic resources in molluscan aquaculture. Rev Aquacult 1, 251–259.

Hedgecock, D., Mcgoldrick, D.J., Bayne, B.L., 1995. Hybrid vigor in Pacific oysters: an experimental approach using crosses among inbred lines. Aquaculture 137, 285–298.

Hochholdinger, F., Hoecker, N., 2007. Towards the molecular basis of heterosis. Trends Plant Sci 12, 427–432.

Ininda, J., Giruchu, L., Njuguna, J., Lorroki, P., 2006. Performance of Three-way cross hybrids for agronomic traits and resistance to maize streak virus disease in Kenya. African Crop Science Journal 14, 287–296.

Jie, T., Gaoyou, Y., Zhigang, L., Fushaomei, L., Yajin, T., University, G.O., 2018. SSR Information Analysis and Unigene Functional Annotation of Transcriptome from *Argopecten irradians* “Bohaihong” and *Argopecten irradians concentricus*. Genomics and Applied Biology.

Kong, L., Song, S., Li, Q., 2017. The effect of interstrain hybridization on the production performance in the Pacific oyster *Crassostrea gigas*. Aquaculture 472, 44–49.

Lautert, L., Lautert, L., Coroilli, E.M., Witt, R.R., 2012. Reproductive isolation of inter-specific crosses between *Argopecten irradians* and *Argopecten purpuratus*. Marine Sciences 36, 9–14.

Liang, S., Luo, X., You, W., Luo, L., Ke, C., 2014. The role of hybridization in improving the immune response and thermal tolerance of abalone. Fish Shellfish Immun 39, 69–77.

Liu, Z., 2007. Upper incipient lethal temperature of *Argopecten irradians concentricus* Say. Journal of Fishery Sciences of China.

Nan, C., Xuan, L., Gu, Y., Han, G., Dong, Y., You, W., Ke, C., 2016. Assessment of the thermal tolerance of abalone based on cardiac performance in *Haliotis discus hannai*, *H. gigantea* and their interspecific hybrid. Aquaculture 465, 258–264.

Nan, L., Zhang, J., Feng, W., Liu, X., Li, Z., Wang, C., 2012. Inter-specific Hybridization Between *Argopecten purpuratus* and *Argopecten irradians concentricus*. Chinese Agricultural Science Bulletin 28, 131–135.

Pan, Y., Jie, F., Xing, L., Wang, W., Ying, P., 2017. Reciprocal crosses between *Argopecten irradians concentricus* and *Chlamys nobilis*. Journal of Fishery Sciences of China 24, 698–709.

Saleh, G.B., Abdullah, D., Anuar, A.R., 2002. Performance, heterosis and heritability in selected tropical maize single, double and three-way cross hybrids. J Agr Sci-Cambridge 138, 21–28.

Samain, J., McCombie, H., Samain, J.F., 2008. Summer Mortality of Pacific Oyster *Crasostrea Gigas*. Quae.

Stelkens, R.B., Schmid, C., Selz, O., Seehausen, O., 2009. Phenotypic novelty in experimental hybrids is predicted by the genetic distance between species of cichlid fish. Bmc Evol Biol 9, 283.

Stone, D.A.J., Bansemer, M.S., Lange, B., Schaefer, E.N., Howarth, G.S., Harris, J.O., 2014. Dietary intervention improves the survival of cultured greenlip abalone (*Haliotis laevigata Donovan*) at high water temperature. Aquaculture 430, 230–240.

Sun, J.Z., Wang, M.X., Chu, C.J., 2004. Experiment on “Penglai Red” scallop culture. Fishery Modernization, 1–5.

Wang, C., Li, Z., 2010. Improvement in production traits by mass spawning type crossbreeding in bay scallops. Aquaculture 299, 51–56.

Wang, C., Liu, B., Liu, X., Ma, B., Zhao, Y., Zhao, X., Liu, F., Liu, G., 2017. Selection of a new scallop strain, the Bohai Red, from the hybrid between the bay scallop and the Peruvian scallop. Aquaculture 479, 250–255.

Wang, C., Liu, B., Li, J., Liu, S., Li, J., Hu, L., Fan, X., Du, H., Fang, H., 2011. Introduction of the Peruvian scallop and its hybridization with the bay scallop in China. Aquaculture 310, 380–387.

Wang, H., Liu, J., Li, Y., Zhu, X., Liu, Z., 2014. Responses to two-way selection on growth in mass-spawned F1 progeny of *Argopecten irradians concentricus* (Say). Chin J Oceanol Limn 32, 349–357.

Wang, L.L., Zhang, H., Song, L.S., Guo, X.M., 2007. Loss of allele diversity in introduced populations of the hermaphroditic bay scallop *Argopecten irradians*.. Aquaculture 271, 252–259.

Wilbur, A.E., Gaffney, P.M., 1997. A genetic basis for geographic variation in shell morphology in the bay scallop, *Argopecten irradians*. Mar Biol 128, 97–105.

Xu, H., Li, Q., Kong, L., Yu, H., Liu, S., 2019. Fertilization, survival and growth of hybrids between Crassostrea gigas and Crassostrea sikamea. Fisheries Sci, 1–8.

Yamak, U.S., Sarica, M., Boz, M.A., 2014. Comparing slow-growing chickens produced by two-and three-way crossings with commercial genotypes. 1. Growth and carcass traits. Eur Poultry Sci 78.

Yan, X., Zhang, Y., Sun, H., Huo, Z., Sun, X., Yang, F., Zhang, G., 2010. Three way crosses between two-band red and white zebra strains of Manila clam, *Ruditapes philippinarum*. Journal of Fisheries of China 34, 1190–1197.

Yang, A., Wang, Q., Liu, Z., Zhou, L., 2004. The hybrid between the scallops *Chlamys farreri* and *Patinopecten yessoensis* and the inheritance characteristics of its first filial generation. Marine Fisheries Research 25, 1–5.

You, W., Ke, C., Luo, X., Wang, D., 2009. Growth and survival of three small abaloneHaliotis diversicolor populations and their reciprocal crosses. Aquac Res 40, 1474–1480.

You, W., Guo, Q., Fan, F., Ren, P., Luo, X., Ke, C., 2015. Experimental hybridization and genetic identification of Pacific abalone *Haliotis discus* hannai and green abalone *H*. *fulgens*. Aquaculture 448, 243–249.

Zhang, F., He, Y., Yang, H., 2000. Introduction engineering of bay scallop and its comprehensive effects. Engineeringence 2, 30–35.

Zhang, H., Liu, X., Zhang, G., Wang, C., 2007. Growth and survival of reciprocal crosses between two bay scallops, *Argopecten irradians concentricus* Say and *A*. *irradians irradians Lamarck*. Aquaculture 272, S88–S93.

Zhang, S., Li, L., Wu, F., Zhang, G., 2014. Yield trait improvement of bay scallops following complete diallel crosses between different scallop stocks. Chin J Oceanol Limn 32, 1–7.

Zhang, Y., Zhang, Y., Li, J., Xiao, S., Xiang, Z., Wang, Z., Yan, X., Yu, Z., 2016. Artificial interspecific backcrosses between the hybrid of female *Crassostrea hongkongensis* × male *C*. *gigas* and the two parental species. Aquaculture 450, 95–101.

Zhang, Z., Chen, J., Li, L., Tao, M., Zhang, C., Qin, Q., Xiao, J., Liu, Y., Liu, S., 2014. Research advances in animal distant hybridization. Science China Life Sciences 57, 889–902.

Zheng, H., Xu, F., Zhang, G., 2011. Crosses between two subspecies of bay scallop *Argopecten irradians* and heterosis for yield traits at harvest. Aquac Res 42, 602–612.

Zheng, H., Zhang, G., Guo, X., Liu, X., 2008. Inbreeding depression for various traits in two cultured populations of the American bay scallop, *Argopecten irradians irradians* Lamarck (1819) introduced into China. Journal of Experimental Marine Biology & Ecology 364, 42–47.

